# Activity flow underlying abnormalities in brain activations and cognition in schizophrenia

**DOI:** 10.1101/2020.12.16.423109

**Authors:** Luke J. Hearne, Ravi D. Mill, Brian P. Keane, Grega Repovš, Alan Anticevic, Michael W. Cole

**Author notes:** **Corresponding Author:** Luke Hearne, Center for Molecular and Behavioral Neuroscience, Rutgers University, 197 University Avenue, Newark, NJ 07102.

## Abstract

Cognitive dysfunction is a core feature of many brain disorders such as schizophrenia (SZ), and has been linked to both aberrant brain functional connectivity (FC) and aberrant cognitive brain activations. We propose that aberrant network activity flow over FC pathways leads to altered cognitive activations that produce cognitive dysfunction in SZ. We tested this hypothesis using activity flow mapping – an approach that models the movement of task-related activity between brain regions as a function of FC. Using fMRI data from SZ individuals and healthy controls during a working memory task, we found that activity flow models accurately predict aberrant cognitive activations across multiple brain networks. Within the same framework, we simulated a connectivity-based clinical intervention, predicting specific treatments that normalized brain activations and behavior in independent patients. Our results suggest that dysfunctional task-evoked activity flow is a large-scale network mechanism contributing to the emergence of cognitive dysfunction in SZ.

## Introduction

Generalized cognitive impairment is one of the most pervasive and stable markers of schizophrenia (SZ) (Kahn and Keefe, 2013; Schaefer et al., 2013). Modern brain imaging techniques, such as fMRI, have linked cognitive dysfunction in SZ to abnormal localized brain activity (Fornito et al., 2012; Pettersson-Yeo et al., 2011). For example, during working memory tasks, individuals with SZ tend to show differences in frontoparietal and default-mode activation compared to healthy controls (Anticevic et al., 2012; Cannon et al., 2005). However, it is likely that cognitive dysfunction emerges in SZ due to abnormal *interactions* between brain regions, not localized activations. This is known as the ‘dysconnection hypothesis’ (Friston et al., 2016; Weinberger, 1993). It is currently unclear how behavioral impairment emerges from the interaction of ‘dysconnected’ FC and aberrant task-evoked activations. Here, to bridge this gap, we link these observations (dysfunctional activity and connectivity) using a recently developed framework termed *activity flow mapping* (Cole et al., 2016).

The last three decades of imaging work have firmly established SZ as a disorder of dysconnectivity (van den Heuvel and Fornito, 2014). Functional connectivity (FC) - defined as the statistical dependence between distinct brain regions - has been instrumental in testing the dysconnection hypothesis, which was originally theorized over a century ago (Bleuler, 1950; Kraepelin, 1919). FC strength tends to be reduced in SZ, with evidence of impaired global network organization (Dong et al., 2018; Pettersson-Yeo et al., 2011; van den Heuvel and Fornito, 2014). Moreover, the interplay between salience, frontoparietal and default-mode networks is particularly impacted in SZ (Supekar et al., 2019). Current thinking suggests that one mechanism underpinning dysconnection in SZ is abnormal N-methyl-D-aspartate (NMDA) receptor mediated synaptic plasticity (Stephan et al., 2006).

Despite the substantial evidence for dysconnectivity, it remains less clear how FC in SZ leads to abnormal brain activations and cognitive deficits. Inspired by connectionist computational modeling principles (Rumelhart et al., 1986), we recently developed *activity flow mapping*, a modeling approach that can be extended to test how distributed sources contribute to localized brain activity (Cole et al., 2016; Ito et al., 2020a). Within this framework, in the context of fMRI, the strength of FC describes the spread of task activations between brain regions. We have shown that this approach is accurate at predicting held-out task activations in both simulated and empirical data from healthy young adults (Cole et al., 2020, 2016; Ito et al., 2017). Critically, applying this method to clinical data allows us to investigate how dysconnectivity and dysfunctional activity flows influence abnormal activations directly tied to deficits in cognition (Mill et al., 2020).

Dysfunctional activity flow could arise in a number of ways. Congruent with the dysconnection hypothesis, it may be that aberrant FC transforms relatively healthy activity in one brain region to dysfunctional activity in another (Mill et al., 2020). Alternatively, relatively healthy FC could propagate preexisting aberrant activations across brain regions. Finally, it may be some mixture of the two, whereby milder ‘subthreshold’ dysfunctional FC interacts with subthreshold aberrant activity to produce suprathreshold dysfunctional activations associated with dysfunctional cognition.

In the current study, we leveraged healthy control (HC, N = 93) and SZ data (N = 36) from the UCLA Consortium for Neuropsychiatric Phenomics LA5c Study (CNP) (Poldrack et al., 2016). Participants completed a spatial capacity working memory (SCAP) task (see **Fig. 1**), which has previously been used to isolate brain activity differences between SZ and HC (Cannon et al., 2005). Using general linear modeling, we first compared task-evoked activations between HC and SZ and identified four differentially activated cortical regions. Then, using activity flow mapping, we tested if these dysfunctional activations in SZ emerged from distributed abnormal activity flows. Finally, within the activity flow framework, we simulated a hypothetical ‘connectivity-based intervention’ to produce new testable hypotheses for improving cognitive deficits in SZ.

**Figure 1.**
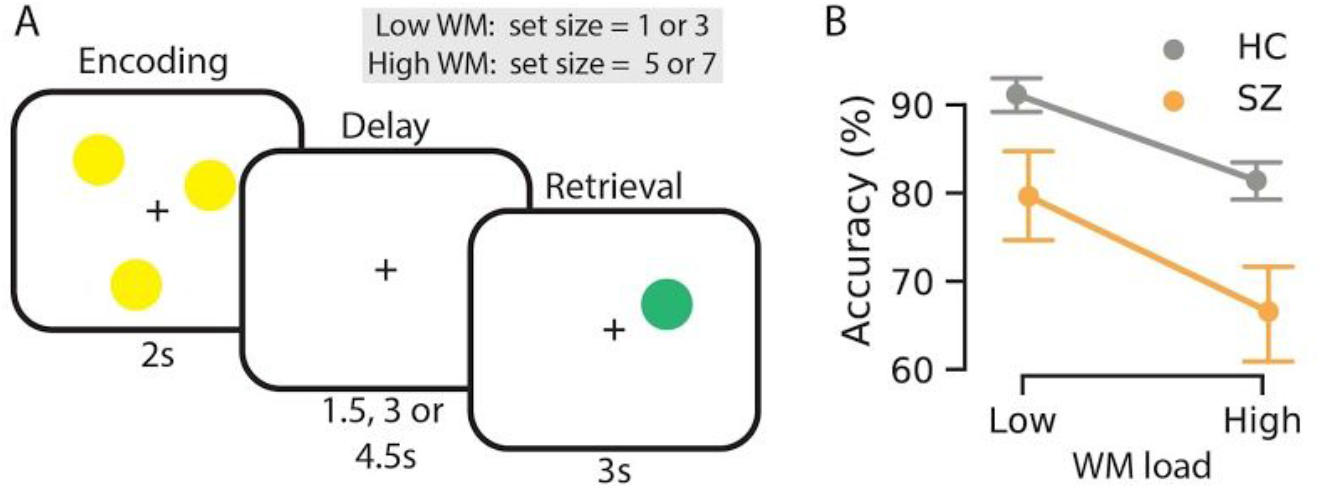
The spatial capacity working memory task (SCAP). **A.** Participants were shown a pseudo-randomly positioned array of 1, 3, 5 or 7 yellow circles. A variable length delay screen is then shown, followed by a single green ‘target’ circle. Participants were asked to indicate whether the green circle was in the same position as any of the yellow circles in the initial array. We contrasted behavior and imaging from the low (set size = 1 or 3) and the high (set size = 5 or 7) working memory conditions. **B.** Task accuracy on the SCAP (chance = 50%). Main effects of working memory and group were both observed.

## Results

### Behavioural differences in spatial working memory

Given that we sought to characterize the brain network mechanisms underlying cognitive dysfunction in SZ, we began by testing for cognitive dysfunction in the SZ group during a spatial working memory task. As expected, participants in the SZ cohort performed less accurately, M_SZ_ = 86.3%, M_HC_ = 73.1%, *t(43.6)* = 4.81, *d* = 1.20, *p* < .001, and slower than the HC group, M_SZ_ = 1237 ms, M_HC_ = 1101 ms, *t(64.9)* = −3.23, *d* = 0.63, *p* = .002. When behavioural accuracy was compared in a two (group: SZ vs. HC) by two (working memory load: low vs. high) mixed ANOVA there were significant main effects of both group [*F(1,127)* = 37.53, η_p_^2^ = .29, *p* < .001] and working memory [*F(1,127)* = 149.5, η_p_^2^ = .54, *p* < .001) (See **Fig. 1B**). However, there was no significant interaction between the two factors [*F(1,127)* = 2.9, η_p_^2^ = 0.02, *p* = .09]. Likewise, when comparing reaction time, main effects of both group [*F(1,127)* = 10.16, η_p_^2^ = 0.07, *p* = .002] and working memory were significant [*F(1,127)* = 259.2, η_p_^2^ = .67, *p* < 0.01]. As above, there was no significant interaction [*F(1,127)* = 1.23, η_p_^2^ = 0.01, *p* = .27]. These results demonstrate that, as expected, the SZ group performed the spatial working memory task worse than the HC group across both low and high WM load conditions.

### Dysfunctional spatial working memory activations in schizophrenia

We next sought to identify localized dysfunctional task-evoked brain activations, which we will subsequently seek to predict via activity flow-related brain network mechanisms hypothesized to underlie cognitive dysfunction in SZ. In response to increased working memory demands both cohorts demonstrated increased activation within dorsal attention and visual networks, and deactivations within the default-mode network (**Fig. 2A**). Four cortical regions were differentially modulated in patients relative to controls (*p* < .05, family wise error [FWE] permutation corrected), demonstrating dysfunctional task-evoked activations. These regions of interest (ROI) included the (i) left ventral anterior cingulate cortex (*ACC*, parcel 57, cingulo-opercular network), (ii) right medial superior temporal area (*MST*, parcel 182, higher order visual network), (iii) right posterior operculum of the sylvian fissure (*PO*, parcel 285, cingulo-opercular network), and (iv) the right posterior insula (*PI*, parcel 347, cingulo-opercular network) (shown by black borders in **Fig. 2A right panel**, parcel borders refer to the original work by Glasser et al., 2016). All four regions were deactivated for high compared to low WM load conditions, and the magnitude of this deactivation was lower for SZ. Likewise, when network-averaged activations were analyzed the default-mode network demonstrated the same pattern of activity with significant differences between groups (**Fig. 2C**, p_FWE_ < .05). Prior work has established reduced deactivations as a hallmark of SZ working memory deficits, and may indicate a lack of spontaneous cognition suppression during working memory task performance (Anticevic et al., 2013, 2012; Landin-Romero et al., 2015).

**Figure 2.**
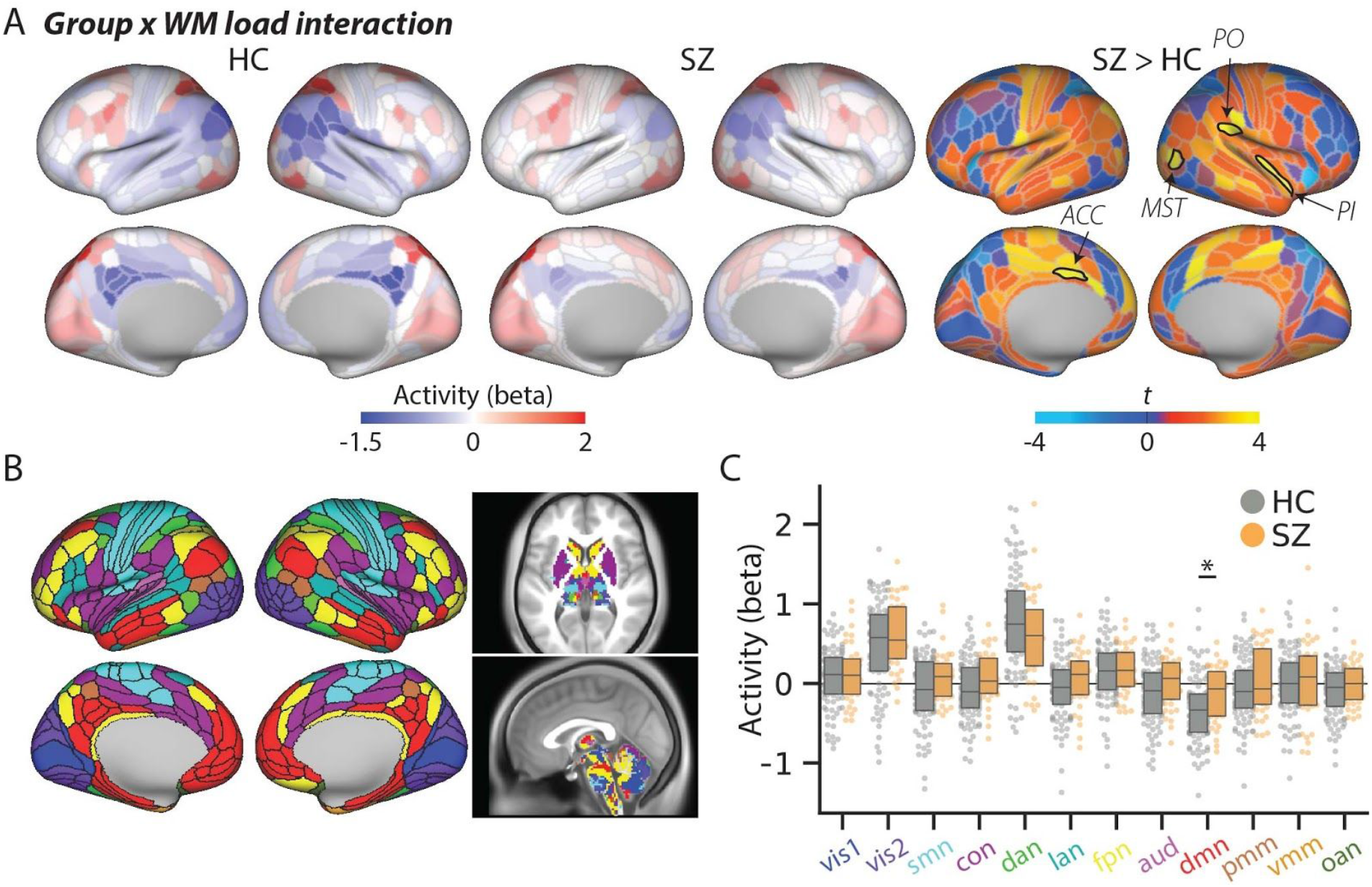
Brain activity associated with the spatial capacity working memory task. **A.** Group average brain activity for the contrast (high > low WM). Significant differences (*p*_FWE_ < .05, 718 comparisons) were found in four cortical regions within the visual and cingulo-opercular network (shown in black borders). **B.** Brain parcellation (718 parcels) and network affiliations (12 networks) used in the study (Ji et al., 2019). No reliable differences were found in the subcortex, therefore visualisation in panel A was limited to the cortex. Subcortical results are presented in fig. S1. **C.** Network level group by working memory brain activation differences. A significant interaction effect was observed within the default-mode network (*p*_FWE_ < .05, 12 comparisons). Network labels (x-axis) match the colors in panel B. Vis1; primary visual, vis2; secondary visual, smn; somatomotor, con; cingulo-opercular, dan; dorsal attention, lan; language, fpn; frontoparietal, aud; auditory, dmn; default-mode, pmm; posterior multimodal, vmm; ventral multimodal, oan; orbito-affective.

In addition to significant activation differences between groups, the activation within each of these brain regions of interest, as well as the default-mode network, correlated with overall task performance (r = −.26 to −.35, *p_bonf_* < .05). Likewise, the average activation across these brain regions correlated with several memory and cognitive control tasks performed outside of the scanner, including measures spanning episodic memory, working memory, fluid reasoning and attention (see **Table S1**). Together these results demonstrate that we identified key dysfunctional cortical regions involved in dysfunctional SZ performance during spatial working memory and broader cognitive demands.

### FC dysconnection in SZ

SZ is considered a disorder of abnormal functional connectivity (Friston et al., 2016; Weinberger, 1993). As such, we tested for group differences in FC between our regions of interest (identified via the task activation analyses reported in the previous section) and the rest of the brain (**Fig. S2**). We found limited differences in FC between groups. In the left MST, we found two differences to/from regions within the right cerebellum (*t*[76.4] = 4.25, *p*_FWE_ = .02) and striatum (*t*[72.6] = 4.01, *p*_FWE_ = .046), such that FC was higher in SZ. We also observed lower FC in SZ between the right PI and the right anterior cingulate cortex (*t*[98.3] = −4.39, *p*_FWE_ = .01). There were no significant FC differences concerning the left ACC and right PO (*p*_FWE_> .05) regions. Moreover, when averaging FC within/across networks we found no statistical differences between groups (*p*_FWE_> .05).

### Activity flow mapping predicts dysfunctional activations in SZ

Inspired by connectionist neural network modeling principles (Rumelhart et al., 1986, Ito et al., 2019), activity flow mapping tests the idea that task-evoked activity is propagated between brain regions via distributed processes captured by FC (Cole et al., 2016). Each held-out ‘target’ activation is modelled as the sum of all other task activation amplitudes weighted by their FC with the target brain region (see **Fig. 3A**). Performed iteratively, activity flow mapping results in a set of brain activity predictions for each region, experimental condition and participant in the dataset. This approach has previously been validated in healthy individuals (Cole et al., 2020, 2016).

**Figure 3.**
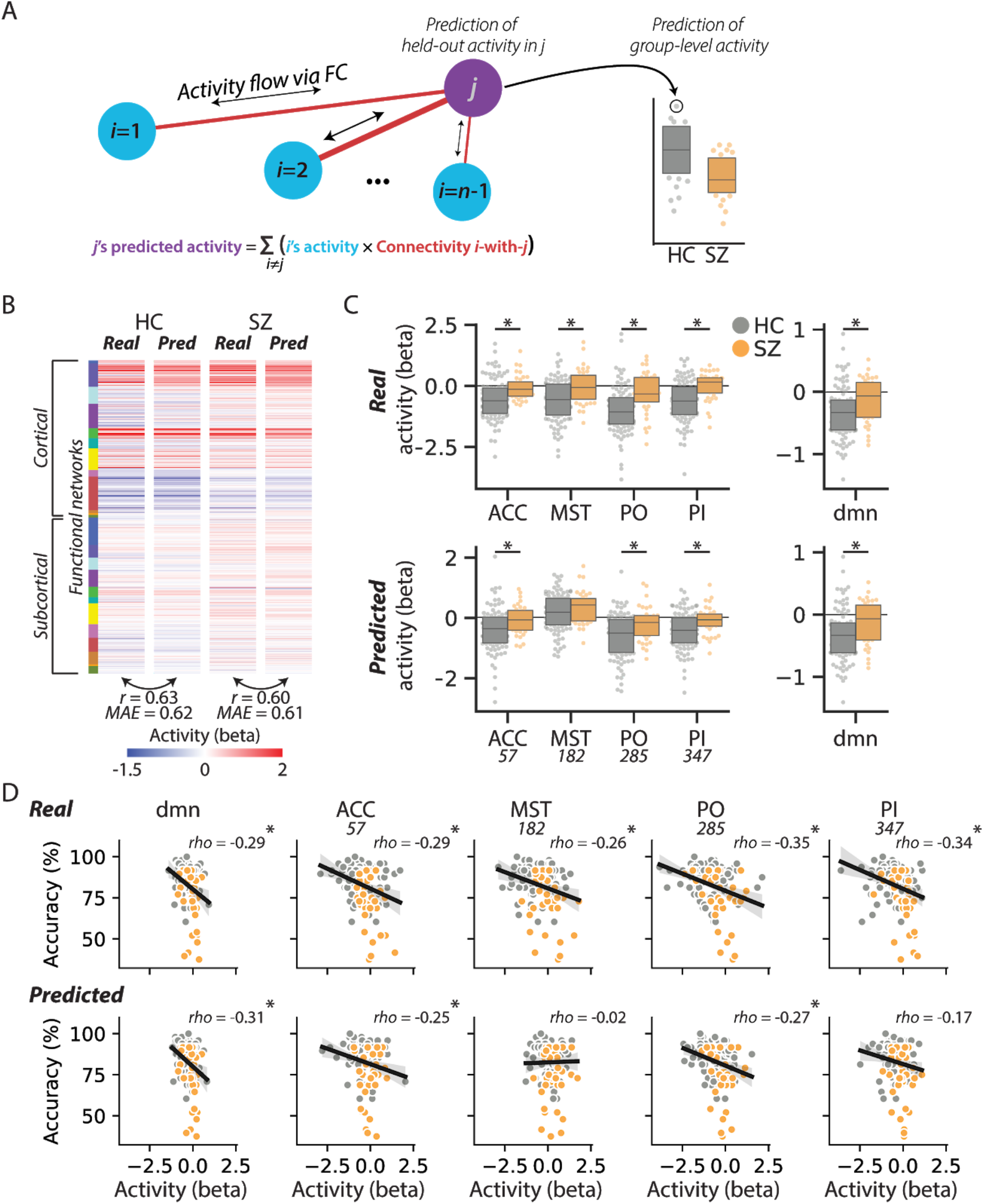
Predicting dysfunctional activity with activity flow mapping. **A.** Schematic of the activity flow algorithm. The task-evoked activation of brain region *j* can be predicted by summing the activity of all other brain regions (*i*) weighted by their connectivity with *j*. The critical assumption of activity flow is that activations are produced by distributed processes that are captured by FC estimates. **B.** Group averaged empirical and predicted (Pred) activations (high > low WM) for healthy control and schizophrenia groups. Note that *r* and *MAE* statistics were conducted at the participant level and then averaged (group averages are shown visually). **C.** Real (top) and predicted (bottom) activations for the four regions of interest and the default-mode network. Aside from the right MST located in the visual cortex, group differences could be captured in the activity flow predicted data. **D.** Correlations between SCAP accuracy scores (*y*-axis) and real activity (top panel) or predicted brain activity (bottom panel) for each region of interest. * indicates *p* < .05. As noted in text, the exploratory empirical analyses were family-wise error corrected for 718 comparisons whereas the confirmatory analyses were bonferroni corrected for four and five comparisons (panel C and D, respectively).

We tested whether activity flow mapping predictions could recapitulate the network- and region-level brain activity dysfunctions identified in the empirical data (i.e., **Fig. 2**). Activity flow mapping was applied to every subject to predict activity in low and high working memory demand conditions. Activity was then subjected to the same contrast used in the empirical data - low versus high working memory demands - generating a single whole-brain activity vector for each participant. To assess activity flow predictions at the whole-brain level, for each subject the real and predicted data were compared via correlation, mean absolute error (MAE) and the coefficient of determination (R^2^). Critically, the four ROI were held out of the activity flow prediction; this ensured that accurate predictions did not rely upon simply transferring dysfunction from one significant dysfunctional region to another. Repeating the analysis including the four held out regions did not alter the results (see Supplementary material).

Across both groups activity flow mapping successfully predicted activity patterns across the whole brain, *r*_HC_ = .63 (one-sample *t*-test compared to zero, *t*[92] = 57.2, *p* < .001), *r*_SZ_ = .60 (*t*[35] = 31.4, *p*< .001), MAE_HC_ = 0.62, MAE_SZ_= 0.61, R^2^_HC_= .40 (*t*[92] =26.0, *p*< .001), R^2^_SZ_ = .35 (*t*[35] =13.6, *p* < .001) (**Fig. 3B**). When compared, predictions were not significantly better for either group (*p* > .15 across all measures).

Next, we chose to focus on regions that had shown statistically robust group differences in the empirical data (i.e., those in **Fig. 2A**, p_FWE_ corrected < .05). For each of these specific regions we performed a between groups *t*-test on the *predicted* activation data. Group differences were observed in three of the four regions; left ACC: *t*(95.4) = 3.01, *p_bonf_*= .014, right MST: *t*(72.5) = 1.64, *p_bonf_*= .425, right PO: *t*(82.3) = 3.39, *p_bonf_* = .004, right PI: *t*(88.4) = 3.45, *p_bonf_* = .003 (bonferroni corrected for four comparisons). Additionally, these predictions all mirrored the pattern of empirical data whereby healthy controls were characterized by decreased WM load activity relative to SZ. The same pattern of results was found in the default-mode network: *t(74.5)* = 3.05, *p* = .003 (**Fig. 3C**).

### Correlations with individual differences in behavior

We next correlated individual differences in actual and predicted activity with working memory task accuracy. We found all regions of interest were negatively correlated with behavior, such that greater deactivation was related to improved task accuracy (left ACC; *rho* = −.29, *p_bonf_*< .001, right MST; *rho* = −.26, *p_bonf_* = .012, right PO; *rho* = −.35, *p_bonf_*< .001, right PI; *rho* = −.34, *p_bonf_*< .001, dmn; *rho* = −.29, *p_bonf_*= .004, bonferroni corrected for five comparisons) (**Fig. 3D** upper panel). Using activity flow mapping the magnitude and direction of these results could be replicated for most comparisons (left ACC; *rho* = −.25, *p_bonf_*= .023, right PO; *rho* = −.27, *p_bonf_* = .008, dmn; *rho* = −.31, *p_bonf_*= .001), but not for the right MST (*rho* = .02, *p* = .86) or PI (*rho* = −.17, *p_bonf_* = .25, bonferroni corrected for five comparisons, **Fig. 3D** bottom panel).

### Activity flow contributions to dysfunctional activity

Having established that activity flow mapping accurately predicts group-level dysfunction in brain activity, we sought to investigate how such differences arise within the model. Recall that a given activity flow estimate is the *sum of* individual flow terms (*i*’s activity x connectivity *i*-with-*j*). Activity flow terms therefore represent a potential brain-wide map capturing the regional contributions that give rise to a target activation magnitude. Thus, we investigated how these individual flow terms differed across the two groups, giving rise to dysfunctions in activity (**Fig. 4**).

**Figure 4.**
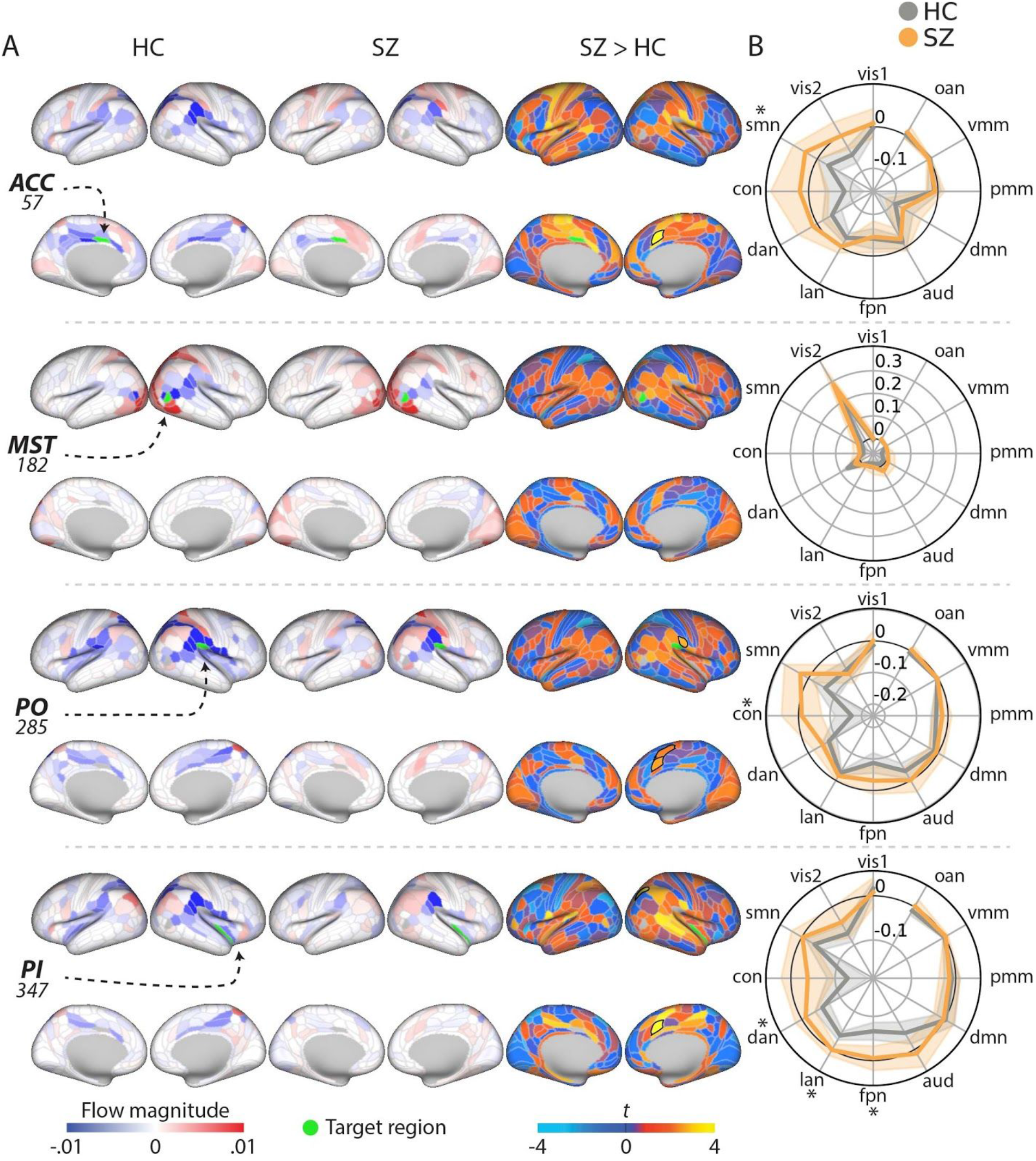
Activity flow contributions to localised dysfunctional activity. **A.** Region-specific activity flow terms (i.e., region *i*’s activity x connectivity *i*-with-*j*) used to predict the target activation (rows) within each cohort. The sum of all terms equal the final activity flow prediction. These spatial maps represent a plausible model of how an individual activation emerges within the activity flow mapping framework. Black borders indicate *p*_FWE_ <.05 (718 comparisons). Subcortical results are presented in sfig.3 **B.** Summary polar plots indicating the summation of activity flow terms within each network. The shaded patches indicate 95% confidence intervals. * indicates *p*_FWE_ < .05 (12 comparisons).

The regions of interest within the cingulo-opercular network (ACC, PO, PI) tended to have a spatially similar activity flow profile when contrasting high and low working memory demand (**Fig. 4A**). This pattern was characterized by negative activity flow from the inferior parietal lobule in HC. In addition to this common pattern, each region had a distinct pattern of activity flow contributions.

We found select differences in activity flow terms between groups when analyzed at the brain region level. The right anterior cingulate cortex showed consistently higher activity flow terms in SZ across the three ROI (ACC; *t*[61.48] = 3.87, *p*_FWE_ = .03, PO; *t*[79.0] = 3.91, *p*_FWE_ = .02, PI; *t*[81.1] = 3.78, *p*_FWE_ = .03). The same pattern of increased SZ activity flow terms were observed in the right PO regarding the right supplementary motor area (*t*[116.4] = 3.86, *p*_FWE_ = .02) and posterior operculum (*t*[69.8] = 3.78, *p*_FWE_ = .03), as well as the right PI and the intraparietal area (*t*[88.3] = 4.20, *p*_FWE_ = .007).

At the network level, for the left ACC and right PO the groups differed in activity flow terms from the sensory-motor [t(88.5) = 2.95, *p_FWE_* = .025] and the cingulo-opercular network [t(86.27) = 3.84, *p_FWE_* = 0.001], respectively (**Fig. 4B**). In the right PI, groups differed across dorsal attention, fronto-parietal and language functional networks [t(75.5) = 2.70, *p_FWE_* = .034, t(83.7) = 4.10, *p*_FWE_ < .001, t(74.7) = 3.61, *p_FWE_ =* .001]. All of these group differences were characterized by increased activity flow terms in the SZ compared to the HC cohorts. Overall, these results suggest dysfunctional activity flow between the source regions and sensorimotor or cognitive control networks in SZ.

As noted in the prior section, activity flow mapping did not produce accurate predictions for the right MST located in the visual cortex. As shown in **Fig. 4** this was due to within-network activity flow terms dominating the predicted values. This is in line with recent evidence suggesting that activity flow mapping is less accurate in regions that are lower in the cortical hierarchy (e.g., in visual cortex) due to distributed activity influencing those regions less (Ito et al., 2020b).

### Simulating functional connectivity changes to normalise patient brain activity and behaviour

In our final analysis we sought to simulate a hypothetical connectivity-based treatment for SZ. Brain stimulation techniques that alter FC are a potential focal treatment option for psychiatric disorders (Cocchi and Zalesky, 2018). Therefore, we extended the activity flow mapping framework to investigate the feasibility of changes in FC resulting in normalized dysfunctional brain activations and cognition. Results from this analysis have the potential to generate testable hypotheses guiding future brain stimulation interventions.

In brief, we used a linear regression model to fit empirical SZ activations to the average healthy activation for each region of interest. The model weights were then used to derive the simulated connectivity intervention for each individual (**Fig. 5A**, see Methods for full details). The difference between the average empirical FC and the simulated FC is shown in **Fig. 5B**. Overall, our data-driven connectivity intervention demonstrated increased FC between each target region and regions within the parietal and prefrontal cortices, in conjunction with decreased sensory network (visual and motor cortices) FC would serve to normalize dysfunctional activations and behavior. The four simulated interventions were highly correlated with each other (*r*_mean_ = .71), suggesting a single connectivity intervention might normalize activity for all four regions. Moreover, the connectivity intervention decreased the similarity in group-averaged FC between the SZ and HC (r_mean_ = .54), compared to the empirical data (r_mean_ = .92). This suggests that the regression model did not simply replace the existing SZ FC weights with those more similar to healthy participants.

**Figure 5.**
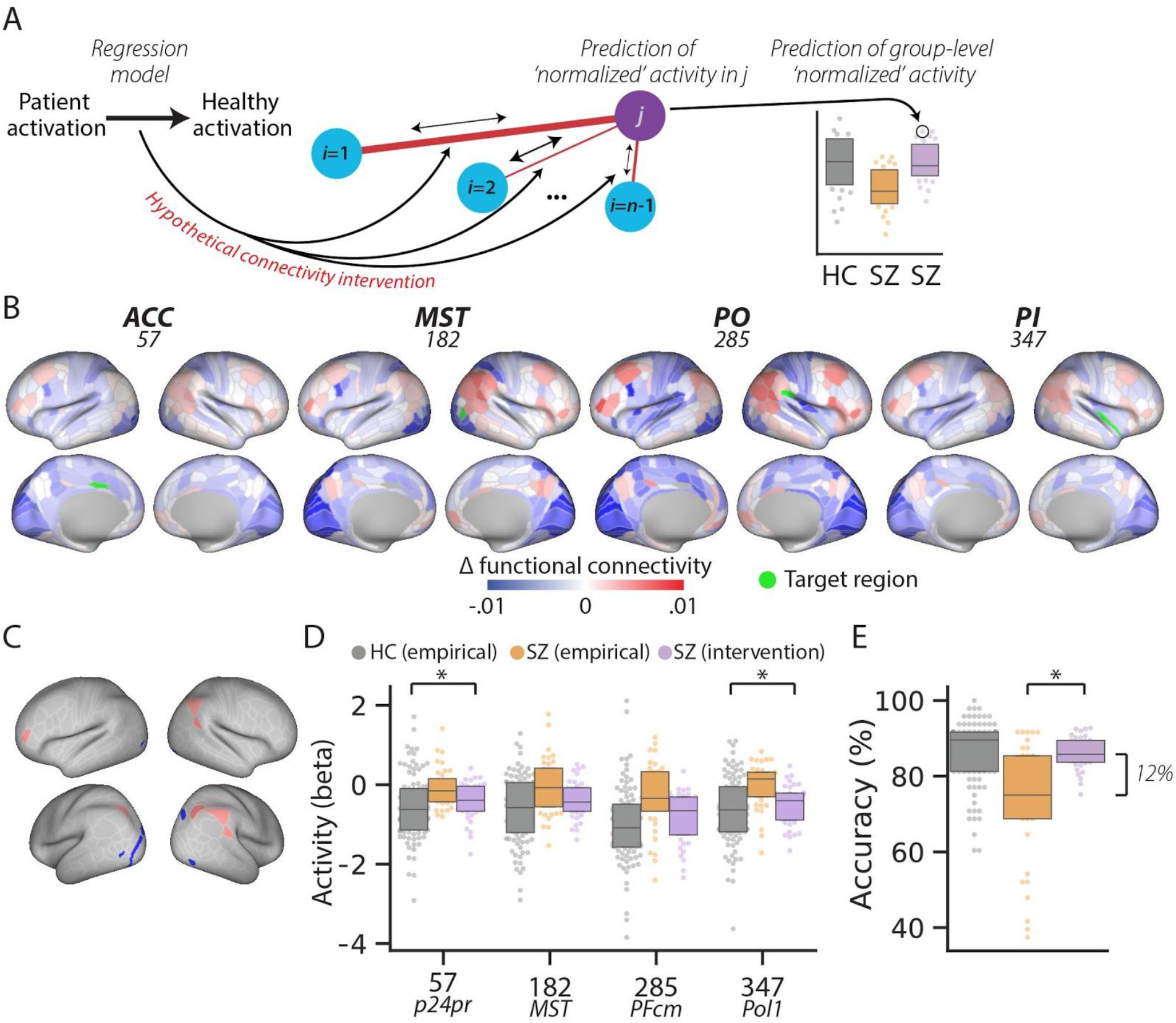
Hypothetical connectivity intervention in schizophrenia. **A.** A regression model was used to optimize SZ functional connectivity to best fit the HC data. The simulated FC was then used to predict activations in the activity flow framework. **B.** Average data-driven FC ‘intervention’ weights for each region of interest in the SZ cohort. The four simulated interventions were highly correlated with each other (*r*_mean_ = .92), despite the interventions being calculated independently for each target region. **C.** The top/bottom five cortical regions requiring the largest connectivity intervention across regions of interest **D.** The FC intervention was verified by applying activity flow mapping with the altered FC; SZ activation levels were normalised (purple) compared to empirical SZ activity (orange) and HC activity (grey). Importantly, the interventions were calculated and tested using cross-validation, with separate subjects used for intervention calculation and testing. **E.** Predicted behavior generated from simulated FC (purple) compared to the empirical task accuracy. By altering SZ functional connectivity our model suggests that behavior would be improved.

To verify the simulated FC treatment we repeated the original activity flow mapping analysis to predict a new set of task-evoked activations. We compared the empirical SZ activity values to the predicted activations in SZ (**Fig. 5D**). For two of the four regions, the predicted activations significant differed in the same direction as the HC empirical data; left ACC: *t*(70) = 2.96, *p_bonf_*= .02 (corrected for four comparisons), right PI: *t*(70) = 3.62, *p_bonf_*= .002. For the other two regions, activity was decreased but not significantly; right PO: *t*(70) = 2.34, *p_bonf_*= .08, right MST: *t*(70) = 2.07, *p_bonf_*= .17). Then, leveraging the existing relationship between the empirical activations and behavioral task accuracy (i.e., **Fig. 4D**), we fit a support vector regression model with activations predicting behaviour and applied the model to the newly altered activations (see **Methods**). This process resulted in a new predicted task accuracy for each individual in the SZ cohort (**Fig. 5E**). The predicted task accuracies showed marked improvement over the original SZ behavior (12.8% difference), *t*(70) = −4.76, *p* < .001). These results demonstrate the plausibility of connectivity-based SZ treatments resulting in normalized cognitive activations and improved cognitive function in SZ.

## Discussion

Cognitive impairment is a core feature of SZ and is related to both aberrant FC and abnormal task-evoked activity (van den Heuvel and Fornito, 2014). In line with the ‘dysconnection’ hypothesis (Friston et al., 2016), we proposed that aberrant network interactions (activity flows) lead to altered cognitive activations that produce dysfunctional behavior. To test this prediction, we used activity flow mapping to model the movement of task-related activity between brain regions as a function of FC. We showed that behavior-related dysfunctional activations could be accurately predicted from spatially distributed sources, suggesting that FC plays a key role in producing aberrant activity and behavior in SZ. Specifically, we observed increased activity flow between sensorimotor and cognitive control networks in SZ, which resulted in clinically-relevant *reduced* deactivations suggestive of an inability to deactivate distracting information. Building on these results, using data-driven simulations, we found that altering regions within the parietal and prefrontal cortices provided the most optimal intervention in normalizing activity and behavioural performance in individuals with SZ.

Deficits in working memory have been consistently observed in SZ (Heinrichs, 2005; Schaefer et al., 2013). Cognitive tasks that engage working memory typically involve activation of the FPN and deactivation of the DMN. Increased dissociation between these two systems is thought to be beneficial for task performance (Anticevic et al., 2012; Fox et al., 2005; Hearne et al., 2015). While both HC and SZ cohorts engaged these networks, we observed significantly reduced deactivations in SZ within the DMN. Task-evoked deactivations have been interpreted as the suppression of goal-irrelevant functions supported by the DMN (e.g., mind wandering) (Anticevic et al., 2013, 2012) and may be a critical trait marker in SZ (Landin-Romero et al., 2015).

We found the same pattern of reduced deactivation as we observed in the DMN within four cortical areas belonging to the CON and visual network (labelled in **Fig. 2A**). Activation patterns in these regions correlated with performance on the spatial working memory task, as well as other, more general cognitive deficits (e.g., in reasoning and attention). These empirical observations add to a growing literature implicating CON dysfunction in SZ (Dong et al., 2018), which may represent the breakdown in perception-action cycles often observed in SZ (Palaniyappan and Liddle, 2012).

To test the idea that aberrant task-evoked activations emerge from distributed FC, we used activity flow mapping, which is a recently-developed method based on neural network simulations (Cole et al., 2016; Ito et al., 2020a). This approach models a given activation as the FC-weighted sum of all other brain regions’ activity. As in previous work with empirical fMRI data from healthy controls (Cole et al., 2020, 2016) and patients with Alzheimer’s disease (Mill et al., 2020), we found this approach was highly accurate at predicting task-evoked activity across the whole brain at the individual subject level. Indeed, confirming our hypothesis, group differences in activation within the regions of interest were recapitulated by activity flow predictions, suggesting that distributed activity flows over FC play an important role in shaping abnormal task-evoked activation magnitudes in SZ.

Relatively small differences in FC between groups were observed (i.e., only three of 2868 possible connections with the abnormally-activated regions survived multiple comparison correction). This would suggest that dysconnection is unlikely to fully explain the aberrant activations. On the other hand, given that all regions with significantly altered activations were held out of each activity flow modeling analysis, normal FC spreading dysfunctional activity between brain regions is not likely either. Rather, it is likely that both subthreshold dysfunctional FC and activity interact to produce dysfunction activation. Another prominent possibility is that activity flow processes - which are weighted sums of distributed activations - pool a large number of subthreshold aberrant activations (possibly over healthy FC) to produce significant aberrant activations. While large-scale FC dysconnectivity is well characterized in SZ (Dong et al., 2018; van den Heuvel and Fornito, 2014), these results highlight the likely contribution of abnormal local (within-region) processing as well (Shaw et al., 2020). The high accuracy of most of the activity flow predictions suggests that even if diffuse local (within-region) processing is the fundamental cause of SZ dysfunction (e.g., from subtle glutamate receptor malfunctions), that dysfunction is spread and likely pooled via activity flow processes to create significant dysfunctional localized activations.

For each aberrant brain region we examined, the sources of activity flow contributions differed. This result supports the idea that a brain region’s function (or in this case, dysfunction) is determined by its unique connectivity profile (Mars et al., 2018; Passingham et al., 2002). The dysfunctional reduced deactivations observed in SZ were associated with increased activity flow from sensorimotor and cognitive control networks, when compared to HC. However, in the region of interest located within the visual cortex (ROI 182), activity flow mapping performed poorly. This is likely due to the high degree of local processing in that specific brain region, which would not be captured by the assumption of distributed processing within the activity flow framework (Ito et al., 2020b).

Brain stimulation techniques that alter FC are being increasingly seen as a potential focal treatment option for psychiatric disorders (Cocchi and Zalesky, 2018). To gain insight into FC-based treatment in SZ, we simulated a hypothetical connectivity-based intervention. Our simulation suggested that increased FC between the dysfunctional regions of interest and select brain regions in the prefrontal and parietal cortices FPN led to predictions of significantly improved brain activity and behavior. The simulated FC interventions were numerically small, supporting the idea that subtle (though perhaps widespread) changes in FC can have a large impact on behavior (Cole et al., 2014; Krienen et al., 2014) and clinical status (Spronk et al., 2018). Critically, the FC generated by the intervention was less similar to HC than the empirical data, suggesting that simply normalizing the FC was not effective at transforming unhealthy activations. Instead, this would suggest that FC interventions should aim to correct both FC dysfunction, as well as existing abnormal local activity.

Existing attempts to use brain stimulation as a therapeutic intervention in SZ have largely focussed on stimulating DLPFC with mixed outcomes (Kumar et al., 2020; Lett et al., 2014). The evidence for PFC stimulation sites in SZ is supported by its abnormal activation during cognitive control (Callicott et al., 2003; Cannon et al., 2005), its disrupted connectivity profile (Fornito et al., 2011; Meyer-Lindenberg et al., 2001) and neurotransmitter regulation (Lewis and Moghaddam, 2006). Our data-driven simulation complements these observations by corroborating the role of PFC in SZ dysfunction and providing new hypotheses to test regarding particular parietal and temporal lobe regions (see **Fig. 5**). A key avenue for future research will be incorporating data-driven brain models into personalized stimulation treatments (Cocchi and Zalesky, 2018).

We deliberately investigated SZ in a case control design for two reasons. First, the spatial working memory task used here has previously demonstrated clinically relevant group differences in brain activity (Cannon et al., 2005). Second, SZ research has identified abnormal connectivity as a key factor in producing abnormal brain activity and behaviour (Friston et al., 2016; Kraepelin, 1919). However, it is becoming increasingly recognized that psychiatric disorder categories may not carve nature at its joints, resulting in high heterogeneity within disorders, and overlap between disorders (Insel et al., 2010). This is exemplified by recent studies that have demonstrated commonalities in connectivity disruptions across multiple disorders (Sha et al., 2018). Pertinent to the current study, cognitive deficits are also common in many other psychiatric disorders (Diamond, 2013). This suggests that the current results may not be specific to SZ per se, but may reflect general effects observable across multiple disorders.

In conclusion, by linking FC and brain activity in a single methodological approach, we have demonstrated that clinically-relevant activations and behavior in SZ are related to (and plausibly caused by) dysfunctional flow of activity across FC networks. The current results also generate new hypotheses regarding brain stimulation sites for the treatment of cognitive deficits in SZ. Future work should aim to extend the activity flow mapping framework across multiple psychiatric disorders with the aim of developing clinically useful personalized brain models.

## Materials and Methods

### Participants

The data used in this study was obtained from the UCLA Consortium for Neuropsychiatric Phenomics LA5c Study (CNP) via the OpenNeuro database (accession number: ds000030) (Gorgolewski et al., 2017; Poldrack et al., 2016). The CNP contains multimodal brain imaging and behavioural data from healthy adults (n=130) and those with ADHD (n=43), bipolar (n=49) or schizophrenia (n=50) diagnoses. All participants were right-handed. Diagnoses were based on the Structured Clinical Interview for DSM-IV and followed the Diagnostic and Statistical Manual of Mental Disorders, Fourth Edition-Text Revision 10. Full details regarding the original participant recruitment, exclusions and study procedures can be found in the corresponding data paper (Poldrack et al., 2016). Participants gave written informed consent following procedures approved by the Institutional Review Boards at UCLA and the Los Angeles County Department of Mental Health.

For the purposes of the current study we leveraged an age- and sex-matched subset of the healthy control (HC, n = 93) and schizophrenia (SZ, n = 36, exclusions due to missing data and head motion, clarified in subsequent sections) cohorts (see **Table 1** for basic demographics). The majority of participants (n = 27) in the SZ cohort had a schizophrenia diagnosis (DSM-IV-TR), the remaining were diagnosed with schizoaffective disorder (n = 9). Almost all patients at the time of testing were medicated (n = 32, see **S Table 2**).

**Table 1.** Demographics and basic cognitive and clinical measures.

|  | HC (n = 93) | SZ (n = 36) | *p* |
| --- | --- | --- | --- |
| Age, years, mean (SD) | 33 (8.68) | 35.5 (8.87) | 0.16 |
| Sex, n male (%) | 59 (63.44%) | 26 (72.22%) | 0.34 |
| MRI site one, n (%) | 72 (77.42%) | 17 (47.22%) | 0.002 |
| Education, years, mean (SD) | 15.16 (1.59) | 12.78 (1.4) | < 0.001 |
| Head motion, RMS, mean (SD) | 0.06 (0.03) | 0.08 (0.03) | 0.002 |
| *Cognitive measures* |  |  |  |
| Matrix reasoning | 20.43 (4.36) | 15.78 (4.68) | < 0.001 |
| Letter/ Number sequencing | 21.05 (2.89) | 17.75 (3.59) | < 0.001 |
| Vocabulary | 43.48 (8.66) | 32 (8.99) | < 0.001 |
| *Clinical measures* |  |  |  |
| Brief Psychiatric Rating Scale, average score (SD) |  |  |  |
| Positive symptoms |  | 2.8 (1.14) |  |
| Negative symptoms |  | 1.81 (0.76) |  |
| Mania/ disorganization |  | 1.76 (0.76) |  |
| Depression/ anxiety |  | 2.43 (1.15) |  |

### The spatial capacity working memory task

In the current study we focussed on the spatial capacity working memory (SCAP) task, which has previously been used to identify behavioral and brain activation differences between healthy control and schizophrenia cohorts (Cannon et al., 2005; Glahn et al., 2003). During the SCAP participants are shown an array of 1, 3, 5 or 7 yellow circles positioned pseudo-randomly around a fixation cross (2 s). A variable length delay screen is then shown (1.5, 3 or 4.5 s), followed by a single green ‘target’ circle (3 s fixed response). Participants were asked to indicate whether the green circle was in the same position as any of the yellow circles in the initial array. On half the trials, the green and yellow circles were aligned (true-positive), with the other half being true-negative. In total, 48 trials were completed (12 for each array set size, 4 for each delay length). Prior to completing the SCAP in the scanner, participants underwent a supervised instruction and training period.

In the current study we contrasted brain and behavioral data from the 1 and 3 sized arrays (*low working memory*, 24 trials) versus the 5 and 7 sized arrays *(high working memory*, 24 trials), while ignoring the delay factor. The behavioral data from the SCAP was analyzed by contrasting accuracy and mean reaction time between the high and low working memory conditions. The total accuracy (score out of 48) was also used to correlate brain and behavioral variables. A single healthy control subject was excluded due to poor performance on the task (accuracy = 31%, *z* = −5.32).

### Data acquisition and preprocessing

The CNP dataset (Poldrack et al., 2016) was acquired on one of two 3T Siemens Trio scanners at either the Ahmanson-Lovelace Brain Mapping Center (Siemens version Syngo MR B15) or the Staglin Center for Cognitive Neuroscience at UCLA (Siemens version Syngo MR B17). Functional MRI data were collected using a T2* weighted echo-planar imaging sequence (slice thickness = 4mm, 34 slices, TR = 2s, TE = 30 ms, flip angle = 90°, matrix 64 x 64, FOV = 192mm, oblique slice orientation). Functional data acquisition included a resting-state scan and seven task paradigms. Structural MPRAGE scans were used for image preprocessing (TR = 1.9 s, TE = 2.26 ms, FOV = 250 mm, matrix = 256 × 256, sagittal plane, slice thickness = 1 mm, 176 slices). Data collection was split across two seperate days, the order of which were counterbalanced across participants. Prior to further analysis, several participants were excluded on the basis of poor quality, or missing data, as identified by Gorgolewski et al., (2017). Complete details for the CNP data collection and task paradigms can be found elsewhere (Poldrack et al., 2016).

Functional and anatomical data underwent a standard volumetric preprocessing pipeline using fMRIprep (Esteban et al., 2019, version 1.1.8), a nipype based tool (Gorgolewski et al., 2011). Following fMRIprep, the data were further processed using Ciftify (Dickie et al., 2019). Ciftify facilitates the analysis of legacy datasets (such as the CNP, with no T2 weighted structural images) to adopt aspects of the ‘gold standard’ Human Connectome Project approach (Glasser et al., 2016). Ultimately, this allows the analyses to be conducted within ‘grayordinate’ space, incorporating both surface vertices and subcortical voxels, the advantages of which have been outlined in prior research (Coalson et al., 2018; Dickie et al., 2019; Fischl, 2012; Glasser et al., 2016; Van Essen, 2012). See supplementary details for full details of the fMRIprep and Ciftify pipelines. The grayordinate data were then downsampled into the Cole-Anticevic Brain-wide network partition (CAB-NP), a recent whole-brain cortical and subcortical atlas comprised 718 brain regions across the cortex (n=360) and subcortex (n=358) (Ji et al., 2019).

After downsampling, additional standard preprocessing steps were performed on the parcellated resting-state and task-state fMRI data. For the resting-state data, the first 4 TRs were removed. All data were subjected to de-meaning, de-trending and nuisance regression. The nuisance regression pipeline was based on the empirical tests performed by Ciric and colleagues (2017). Specifically, six primary motion parameters were removed, along with their derivatives, and the quadratics of all regressors (24 motion regressors in total). Physiological noise was modeled based on white matter and ventricle signals using aCompCor (Behzadi et al., 2007) within fMRIprep. Five component signals were used, as well as their derivatives, and the quadratics of all physiological noise regressors (20 physiological noise regressors total).

In addition, for the resting-state data we used relative root mean squared displacement (RMS) to identify high movement frames in the data (> 0.25 mm, Satterthwaite et al. 2013). For each of these data points an additional ‘spike’ regressor was added. We also excluded participants with generally high motion (Parkes et al., 2018); any participant with more than 20% of their data in any given functional run above the high motion cutoff (relative RMS > 0.25) were excluded from the analyses (HC = 6, SZ = 12).

The nuisance regression pipeline was completed immediately prior to FC estimation for the resting-state data. For the task-based analyses the regressors were incorporated into the task design matrix.

### Task activation estimation

For the SCAP task, activations were estimated using a standard general linear model (GLM). For each trial, a single boxcar function was used from the onset of the encoding period to the end of the response period (6.5 - 9.5 s depending on delay condition). For each condition (12; 4 working memory x 3 delay) this was convolved with the canonical SPM hemodynamic response function (Friston et al., 1994) and entered into the GLM, as well as the nuisance regressors. The result was a region (718) by condition (12) matrix of regression coefficients representing activation amplitudes for each participant. For the majority of the analyses these activations were averaged across working memory load and subtracted from one another (high - low). For the main analysis, we performed a between groups t-test (SZ > HC) on this contrast, corrected for multiple comparisons (see Statistical analyses section). We also performed this analysis at the level of networks by averaging and contrasting values within the 12 predefined functional networks in the CAB-NP atlas (Ji et al., 2019). Regions and networks that demonstrated a significant group effect were correlated with behavioral data.

### Functional connectivity estimation

Task-general FC was estimated using both resting-state and data from three remaining tasks performed in the scanner (Balloon Analog Risk, Stop Signal, and Task Switching). This decision was motivated by the relatively few timepoints within the resting-state data relative to the number of regions within the brain parcellation (152 timepoints versus 718 regions), as well as the potential for task-state FC to be a better predictor of individual differences (Elliott et al., 2019; Greene et al., 2018) and activity flow estimates (Cole et al., 2020). For the task data we used finite impulse response (FIR) modeling (9 parameters, equivalent to 18 s) to remove the mean task-evoked activation response for each condition. FIR has recently been shown to reduce both false positive and negative rates in the context of task FC estimates (Cole et al., 2019). The nuisance regressors were also added to the GLM. The residuals for each task were concatenated with the resting-state data into a single time series, which were used to calculate FC.

Principal components regression (PCR) was used to estimate FC. Previous work has determined that multiple regression approaches tend to perform better than Pearson correlation within the activity flow mapping framework by removing indirect connections (Cole et al., 2016, Sanchez-Romero & Cole 2019). We opted for PCA regression (as opposed to multiple regression) due to the similar number of overall timepoints to observations in the current study (811 v. 718), which we have used successfully before in datasets with similar properties (Cole et al., 2016; Ito et al., 2017; Mill et al., 2020). In this analysis, rather than using every other timeseries as a predictor for a given brain region (as in multiple regression), a PCA is conducted to limit the number of predictors in the regression model. The resulting beta values are then projected into the original brain region space (from principal component space) to achieve N_*region*_ - 1 beta coefficients (717) which are used as FC edge weights for a given region. The principal components were calculated independently for each to-be-predicted region. When performed across regions, a region x region (718 x 718) FC matrix was computed for each participant. We chose to use the top 100 components in the PCA regression, however we completed control analyses to ensure this did not significantly affect the activity flow mapping results (see Supplementary material).

For each region of interest identified in the GLM, we performed a between groups t-test (SZ > HC) comparing FC values between the region of interest and all other brain regions, corrected for multiple comparisons (see Statistical analyses section). We also performed this analysis at the level of networks by averaging and contrasting values within the 12 predefined functional networks in the CAB-NP atlas (Ji et al., 2019).

### Activity flow mapping

Activity flow mapping was developed as a method to quantify the relationship between FC and task-evoked activations (Cole et al., 2016). Inspired by connectionist principles (Ito et al., 2020a; Rumelhart et al., 1986), activity flow mapping posits that task-evoked activity is propagated between brain regions via functional connectivity. As such, in any given task state, a *target* activation is modeled as the sum of all other *source* activations during the same task, after each activation is multiplied by connectivity between the target and each source.

**Equation 1. The activity flow algorithm**

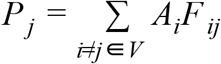

where *P* is the predicted mean activation for region *j* in a given task, *A_i_* is the actual mean activation for region *i* in a given task (a beta value estimated using a general linear model), *i* indexes all brain regions (vector *V*) with the exception of region *j*, and *F_ij_* is the FC estimate between region i and region j. As well as holding out the target region (*j*) from each prediction, any brain region that demonstrated a significant group (SZ vs. HC) task activation effect was also held out. This was to ensure that accurate predictions did not rely upon simply transferring dysfunction from one dysfunctional region to another – rather they had to arise from distributed sources. The algorithm results in a matrix with predicted activations across all nodes and task conditions.

Given a set of predictions that match the original activity data in shape (e.g., region x condition x participant), standard assessments of prediction accuracy, such as those used in machine learning were used (Poldrack et al., 2019). Here we assessed prediction accuracy for each participant using correlation (Pearson *r*), mean absolute error (MAE) and the coefficient of determination (R^2^). Accuracy values were averaged across conditions and participants before being reported in text. Code to conduct activity flow mapping and the subsequent statistics is publicly available via the Brain Activity Flow (“Actflow”) Toolbox (https://colelab.github.io/ActflowToolbox/).

In addition to the standard assessments of accuracy at the participant level noted above, we also tested whether the predicted data could replicate the group-level activity differences observed in the empirical data. To do so, we repeated the high versus low working memory contrast, and group level *t*-tests in the regions of interest (five *t*-tests in total). As in the empirical data, the same regions/networks were correlated with behavior to test whether activity flow predictions preserved behaviorally relevant patterns of activity.

#### Probing activity flow predictions

In the current study we wanted to investigate how dysfunctional activations in SZ arise from distributed activity and connectivity. Assuming activity flow mapping produces accurate predictions, the magnitude of the activity flow terms (i.e., *A_i_F_ij_* in **Equation 1**) represent a plausible model of information/activity-level flow between a given source region and the target region. Group differences in activity flow terms therefore represent dysfunction that is either transferred from a source region (or network) to the target activation, or dysfunction that arises in the target region due to a connectivity-based transformation from source to target. To quantify this, for each dysfunctional region (identified in the GLM) we compared each activity flow term in a between groups *t*-test, corrected for multiple comparisons. We also performed this analysis at the level of networks by summing and contrasting values within the 12 predefined functional networks in the CAB-NP atlas (Ji et al., 2019).

### Simulating a hypothetical connectivity intervention

Considering we have a model of how a dysfunctional localised activation emerges in schizophrenia, an interesting question is raised: what would need to change in the SZ data to normalise dysfunctional activity and behavior? In line with the dysconnection hypothesis (Friston et al., 2016), we sought to develop a simulated FC ‘intervention’ to answer this question. In brief, we used a regression model to fit patient activations to healthy activation levels in the ROI identified by the GLM. The resulting beta weights were interpreted as ‘simulated FC’. We then tested the simulated FC by using activity flow mapping to produce new, *altered*, activity predictions. In a final step, we used the altered activations to generate predictions of SCAP task accuracy which were compared to the original empirical data (see Supplementary material for schematic of pipeline).

#### Hypothetical FC model fitting

Using Pytorch (Paszke et al., 2017) we implemented a linear regression model with gradient descent. Gradient descent was used (rather than standard linear regression) so that the regression weights (*β*) could be initialized as the empirical SZ FC, therefore preserving properties of the empirical data. A separate model was performed for each region of interest. For each regression model, the predictors (*X*) were the individual empirical activations from the SZ cohort and the response variable (*y*) was the average HC value for the same brain region. No intercept was included in the model. We used standard model hyperparameters; the optimizer was stochastic gradient descent (SGD), the loss function was mean standard error loss (MSE) and the learning rate was set to 1e-3. The algorithm was repeated 200 times.

A four-fold cross-validation scheme was used (75% of participants used for training, 25% testing) (Varoquaux, 2018). Within each training set, the regression weights were contrasted with the empirical SZ FC to derive a difference score – the magnitude of the FC intervention. This change in FC was then applied to the empirical SZ FC in the held-out test set to create the hypothetically *altered* FC. The result was a set of altered FC weights for each region of interest and participant that yielded the optimal normalization of their activations.

#### Effect of connectivity intervention on activations and task accuracy

As an alternative to reporting the cross-validated model fit, the altered FC was verified by quantifying the extent to which predictions of brain activity and behavior in SZ became more similar to HC. Thus, in each test set the altered FC and empirical activations were subjected to the activity flow mapping framework (described in previous section) to produce altered activations for the SZ cohort. These values were statistically compared to the SZ empirical data to test whether the existing group effect had been normalized. Showing such an effect would be non-trivial, given that the intervention model was trained on data from independent participants (using cross-validation).

To relate the normalized activations to behavior, we used a support vector regression (SVR) model using default parameters in scikit-learn (Pedregosa et al., 2011) (kernel = rbf, gamma = scale, epsilon = 0.01). For the SVR model, the predictors (*X*) were the empirical activations from participants in the four regions of interest and the response variable (*y*) was total accuracy on the SCAP task (only using data from the training set). This model was then applied to the *altered activations* in the test SZ cohort produced by the hypothetical connectivity intervention, resulting in a behavioral prediction for each SZ participant. The predicted behavior was then statistically compared to the empirical behaviour in the SZ cohort.

### Statistical analyses

Due to the differences in group sizes, Welch’s *t*-test (Welch, 1947) was used for group comparisons. Likewise, due to the non-normal distribution of behavioral variables, correlations were conducted using Spearman’s rank correlation. Where noted, we used the MaxT permutation approach (10,000 permutations) to perform family wise error (FWE) multiple comparison correction (Nichols and Holmes, 2002).

## Data and code availability

All code related to analyses in this study will be publicly released on GitHub. All data are publicly available through https://openneuro.org/datasets/ds000030/.

## Acknowledgements

The authors acknowledge support by the US National Institutes of Health under awards R01 AG055556 and R01 MH109520 to MWC. GR was supported by the Slovenian Research Agency grants J7-5553, J3-9264, P5-0110 and P3-0338. The content is solely the responsibility of the authors and does not necessarily represent the official views of any of the funding agencies. The authors acknowledge the Office of Advanced Research Computing (OARC) at Rutgers, The State University of New Jersey for providing access to the Amarel cluster and associated research computing resources that have contributed to the results reported here. The data used in this manuscript was obtained from the OpenfMRI database (https://openfmri.org/dataset/ds000030/).

## Supplementary material

For full transparency we report the autogenerated fMRIprep preprocessing output, with some edits for clarity. Results included in this manuscript come from preprocessing performed using fMRIPprep 1.1.8 (Esteban et al., 2020, 2018)(RRID:SCR_016216), which is based on Nipype 1.1.3 (Gorgolewski et al., 2011, 2017) (RRID:SCR_002502).

### Anatomical preprocessing

The T1-weighted (T1w) image was corrected for intensity non-uniformity (INU) using ‘N4BiasFieldCorrection’ (Tustison et al., 2010) (ANTs 2.2.0), and used as T1w-reference throughout the workflow. The T1w-reference was then skull-stripped using ‘antsBrainExtraction.sh’ (ANTs 2.2.0), using OASIS as target template. Brain surfaces were reconstructed using ‘recon-all’ (Dale et al., 1999) (FreeSurfer 6.0.1, RRID:SCR_001847), and the brain mask estimated previously was refined with a custom variation of the method to reconcile ANTs-derived and FreeSurfer-derived segmentations of the cortical gray-matter of Mindboggle (Klein et al., 2017) (RRID:SCR_002438).

Spatial normalization to the ICBM 152 Nonlinear Asymmetrical template version 2009c (RRID:SCR_008796) was performed through nonlinear registration with ‘antsRegistration’ (Avants et al., 2008) (ANTs 2.2.0, RRID:SCR_004757), using brain-extracted versions of both T1w volume and template. Brain tissue segmentation of cerebrospinal fluid (CSF), white-matter (WM) and gray-matter (GM) was performed on the brain-extracted T1w using ‘fast’ (Zhang et al., 2001) (FSL 5.0.9, RRID:SCR_002823).

### Functional data preprocessing

For each of the 6 BOLD runs found per subject (across all tasks and sessions), the following preprocessing was performed. First, a reference volume and its skull-stripped version were generated using a custom methodology of fMRIPrep. A deformation field to correct for susceptibility distortions was estimated based on fMRIPrep’s fieldmap-less approach.

The deformation field is that resulting from co-registering the BOLD reference to the same-subject T1w-reference with its intensity inverted (Huntenburg, 2014; Wang et al., 2017). Registration is performed with ‘antsRegistration’ (ANTs 2.2.0), and the process regularized by constraining deformation to be nonzero only along the phase-encoding direction, and modulated with an average fieldmap template (Treiber et al., 2016). Based on the estimated susceptibility distortion, an unwarped BOLD reference was calculated for a more accurate co-registration with the anatomical reference.

The BOLD reference was then co-registered to the T1w reference using ‘bbregister’ (FreeSurfer) which implements boundary-based registration (Greve and Fischl, 2009). Co-registration was configured with nine degrees of freedom to account for distortions remaining in the BOLD reference. Head-motion parameters with respect to the BOLD reference (transformation matrices, and six corresponding rotation and translation parameters) are estimated before any spatiotemporal filtering using ‘mcflirt’ (Jenkinson et al., 2002) (FSL 5.0.9). BOLD runs were slice-time corrected using ‘3dTshift’ from AFNI (Cox, 1996) (RRID:SCR_005927). The BOLD time-series (including slice-timing correction when applied) were resampled onto their original, native space by applying a single, composite transform to correct for head-motion and susceptibility distortions. These resampled BOLD time-series will be referred to as ‘preprocessed BOLD in original space’, or just ‘preprocessed BOLD’.

Additionally, a set of physiological regressors were extracted to allow for component-based noise correction (Behzadi et al., 2007) (CompCor). Principal components are estimated after high-pass filtering the preprocessed BOLD time-series (using a discrete cosine filter with 128s cut-off) for the two CompCor variants: temporal (tCompCor) and anatomical (aCompCor). Six tCompCor components are then calculated from the top 5% variable voxels within a mask covering the subcortical regions. This subcortical mask is obtained by heavily eroding the brain mask, which ensures it does not include cortical GM regions. For aCompCor, six components are calculated within the intersection of the aforementioned mask and the union of CSF and WM masks calculated in T 1w space, after their projection to the native space of each functional run (using the inverse BOLD-to-T 1w transformation). The head-motion estimates calculated in the correction step were also placed within the corresponding confounds file.

All resamplings can be performed with a single interpolation step by composing all the pertinent transformations (i.e. head-motion transform matrices, susceptibility distortion correction when available, and co-registrations to anatomical and template spaces). Gridded (volumetric) resamplings were performed using ‘antsApplyTransforms’ (ANTs), configured with Lanczos interpolation to minimize the smoothing effects of other kernels. Non-gridded (surface) resamplings were performed using ‘mri_vol2surf(FreeSurfer).

Many internal operations of *fMRIPrep* use *Nilearn* 0.4.2 (Abraham et al., 2014) (RRID:SCR_001362), mostly within the functional processing workflow. For more details of the pipeline, see the section corresponding to workflows in fMRIPrep’s documentation (https://fmriprep.readthedocs.io/en/latest/workflows.html).

### Surface-based processing

Ciftify (Dickie et al., 2019) was used to transform the fmriprep generated volumetric functional data into gold-standard Human Connectome Project (HCP) (Glasser et al., 2016) surface ‘grayordinate’ space data. fMRI images were mapped to subject specific MNINonLinear-fsaverage_LR32 grayordinates space (Robinson et al., 2018). Cortical surfaces were based on output from FreeSurfers ‘recon-all’ pipeline. As in the HCP, the data were smoothed 2mm full-width half max Gaussian kernel along the cortical surface.

### Control analyses

fMRI data is thought to have 2 mm to 5 mm of spatial smoothing due to vasculature rather than neural activity (Logothetis and Wandell, 2004). This smoothness could potentially bias activity flow estimates by allowing the target activity to ‘leak’ into the source activity. This would introduce some circularity as information from the target would be used to predict the same target. To confirm this wasn’t the case, we repeated the analyses by excluding all parcels with any vertices within 10 mm of each target region from the set of source regions when calculating FC. Activity flow predictions replicated for the whole brain results, *r*_HC_ = .53, one-sample *t*-test compared to zero, *t*(92) = 37.3, *p* < .001, *r_SZ_* = .48, *t*(35) = 17.8, *p* < .001, and the group differences in specific brain areas, ROI ACC; *t*(95.1) = 2.95, *p_bonf_*= .02, ROI MST; *t*(75.2) = 1.33, *p_bonf_* = .75, ROI PO; *t*(84.9) = 3.31, *p_bonf_* = .005, ROI PI; *t*(858) = 3.25, *p_bonf_* = .005. (bonferonni corrected for four multiple comparisons).

As noted in the main text, we held out the dysfunctional regions of interest from the main activity flow mapping analysis. This was to ensure accurate predictions of dysfunction could not be simply attributed to the transfer between the four regions. To verify this had no bearing on the main results we repeated the analysis when including all regions. We found minimal differences with the results reported in the main text; *r*_HC_ = .63, one-sample *t*-test compared to zero, *t*(92) = 57.4, *p* < .001, *r_SZ_* = .60, *t*(35) = 31.41, *p* < .001, and the group differences in specific brain areas, ROI ACC; *t*(95.2) = 3.08, *p_bonf_*= .01, ROI MST; *t*(72.5) = 1.68, *p_bonf_*= .38, ROI PO; *t*(82.5) = 3.46, *p_bonf_* = .004, ROI PI; *t*(88.5) = 3.50, *p_bonf_* = .004 (bonferonni corrected for four multiple comparisons).

The CNP dataset was collected at two different MRI sites. In the current analysis there were significant differences in the ratio of data collected from the two different MRI sites (77% of data collected from site one in SZ, versus 47% in HC, see **Table 1**). To ensure our results weren’t confounded by MRI site we repeated the analyses in the ROI within a subset of the data demonstrating no MRI site differences between groups (*t*-test between groups, *p* = 0.11,64% versus 47%). All of the current SZ subjects were included (N = 36), but 34 HC subjects were excluded (N = 59). Activity flow predictions replicated for the whole brain results, *r*_HC_ = 0.64, *t*(92) = 46.84, *p* < 0.001, *r*_SZ_ = 0.60, *t*(35) = 31.39, *p* < 0.001 and the group differences in specific brain areas, ROI ACC; *t*(93.0) = 2.62, *p* = .04, ROI MST; *t*(81.9) = 0.89, *p*= .99, ROI PO; *t*(86.1) = 3.07, *p*= .01, ROI PI; *t*(90.1) = 3.01, *p*= .01.

We used PCA regression to estimate functional connectivity. We chose to regress 100 components, however the number of components regressed may affect the final FC estimate and activity flow mapping accuracy. Therefore we repeated the analyses with several component numbers ranging from 50 to 300 to demonstrate that the effect on prediction accuracy was minimal (see **SFig. 3**).

## Supplementary figures and tables

**SFigure 1.**
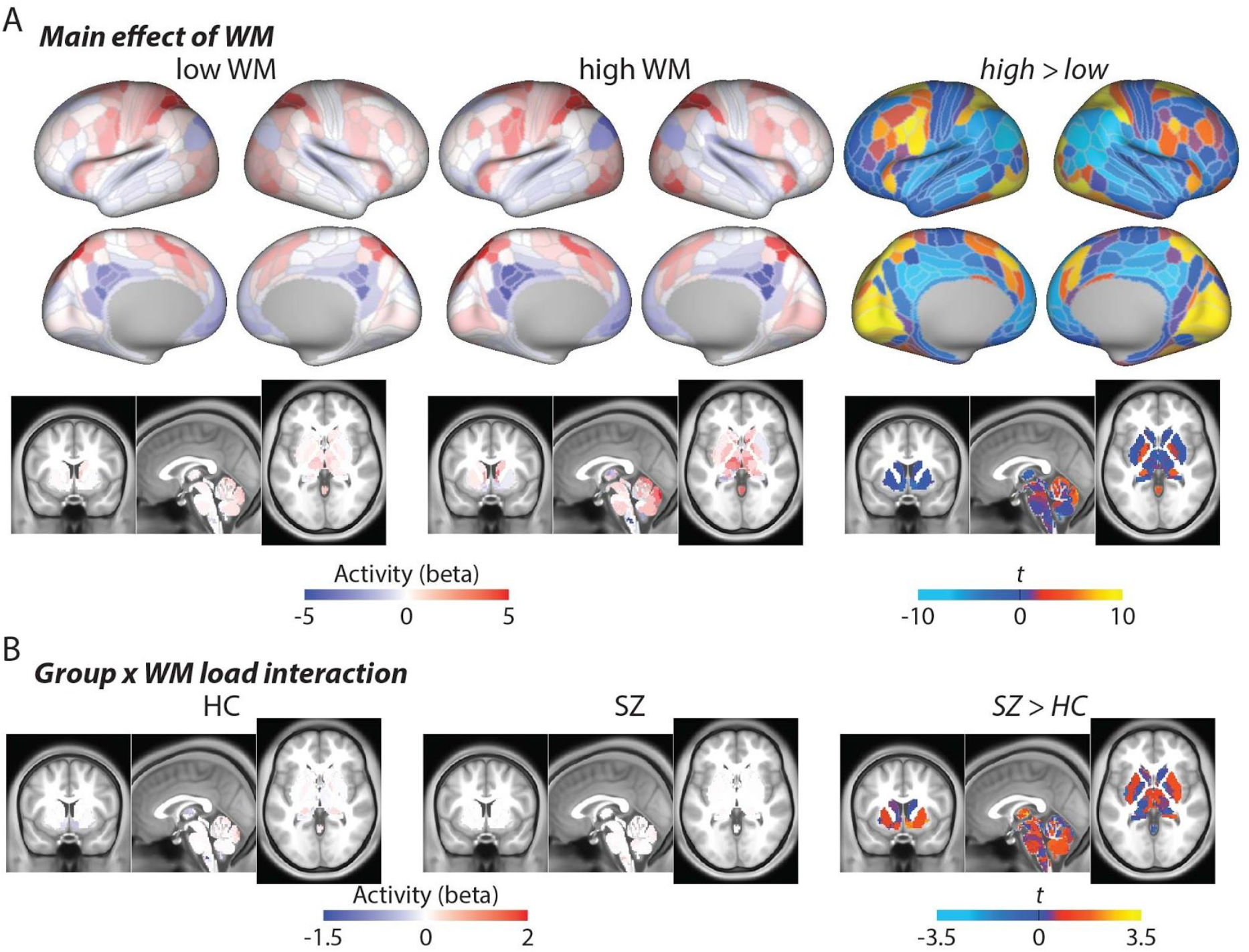
Additional General linear modeling and subcortical results. **A.** Average brain activity (across groups) for low and high working memory conditions. As would be expected, the largest statistical differences were observed in frontoparietal brain regions. **B.** Group average subcortex activity for the contrast (high < low WM). This plot parallels Figure 2A in the main text.

**SFigure 2.**
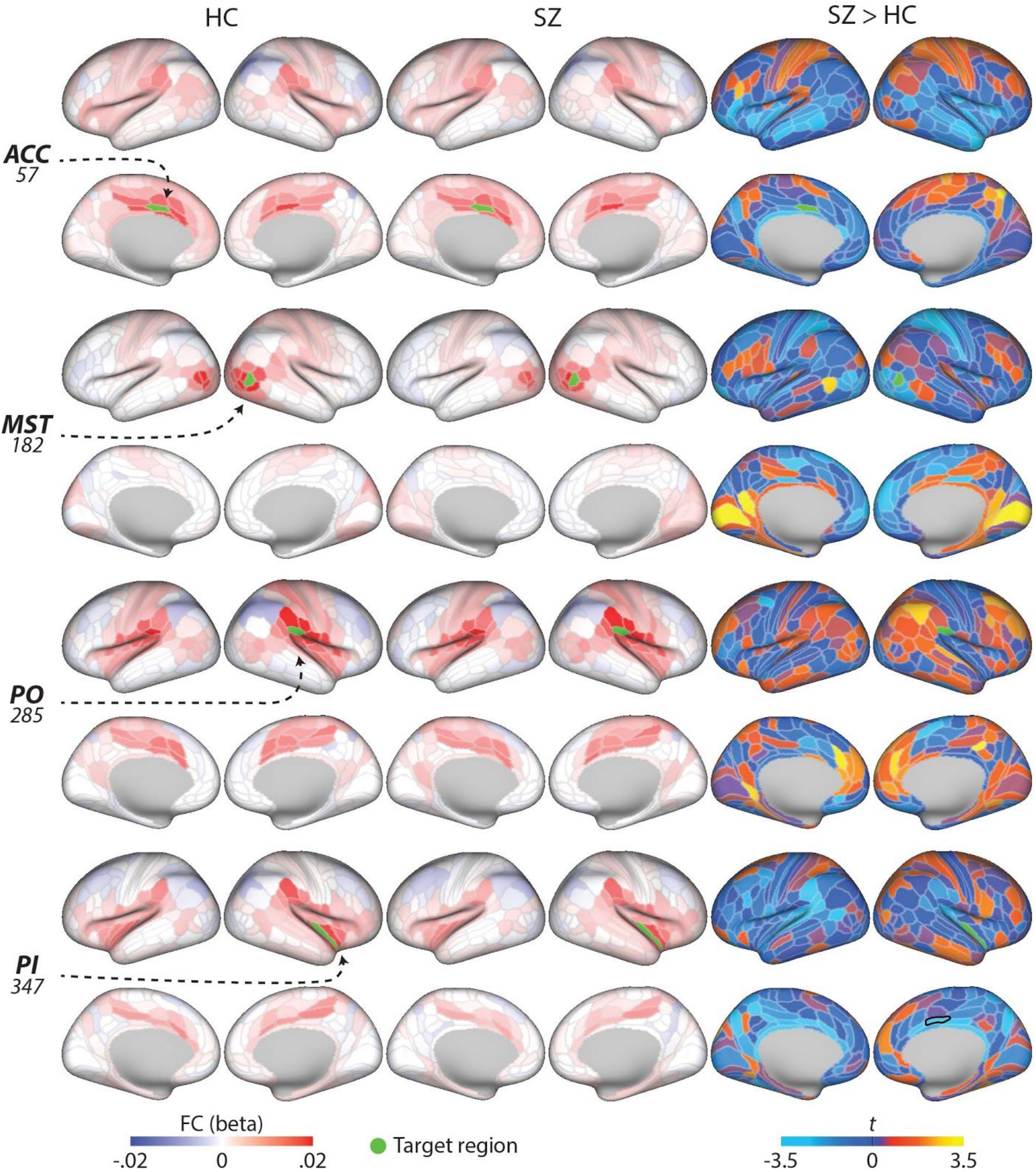
Functional connectivity associated with the four regions of interest. Group average functional connectivity weights plotted for HC (left column) and SZ (middle column) for the four ‘target’ brain regions of interest (highlighted in green). We found limited differences in FC between SZ and healthy controls (*t*-statistic shown in right column). Black borders indicate *p*_FWE_ < .05 (718 comparisons per region).

**SFigure 3.**
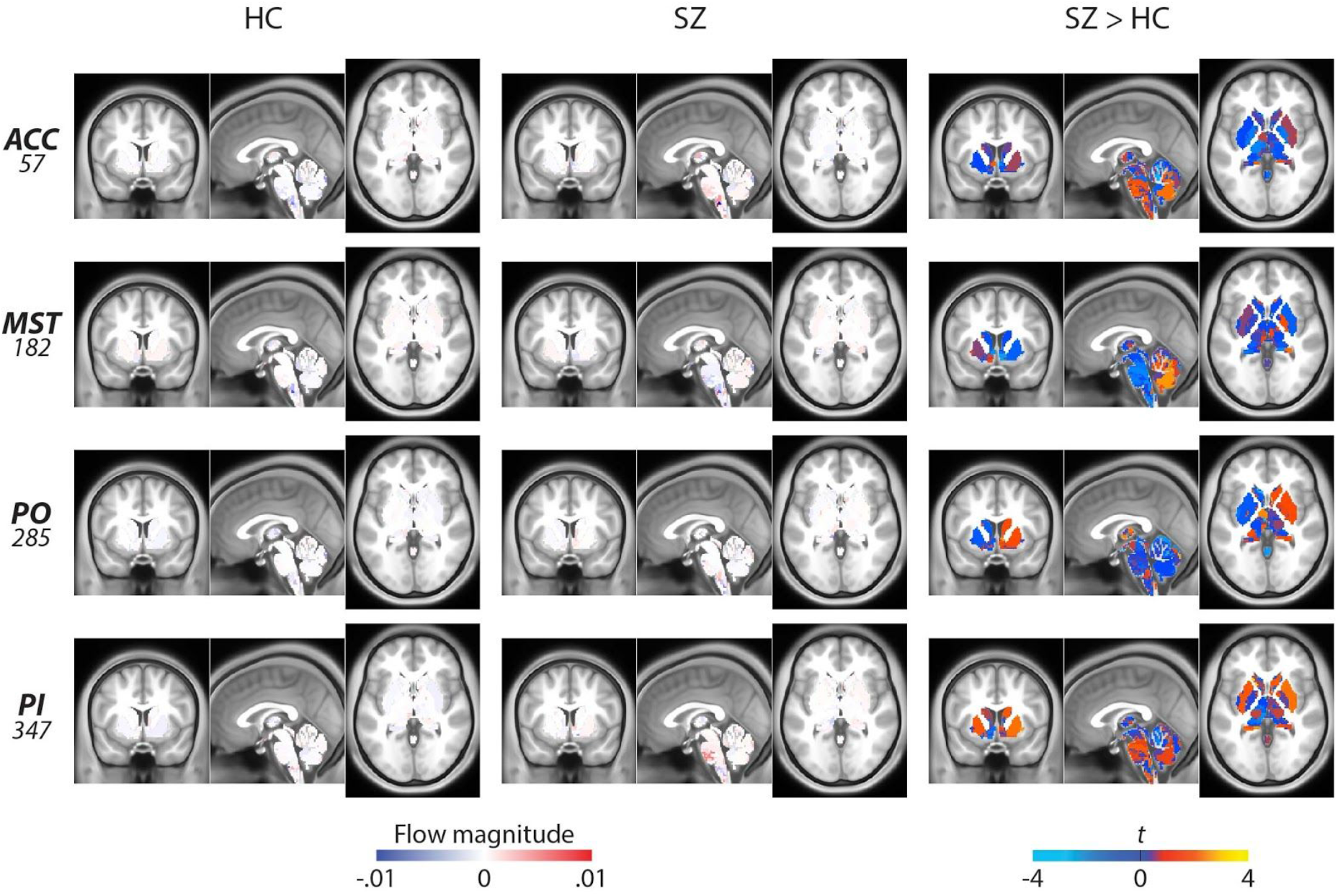
Subcortical activity flow contributions to localised dysfunctional activity. **A.** Region-specific activity flow terms (i.e., region *i*’s activity x connectivity *i*-with-*j*) used to predict the target activation (rows) within each cohort. The sum of all terms equal the final activity flow prediction. These spatial maps represent a plausible model of how an individual activation emerges within the activity flow mapping framework. There were no significant differences in subcortical region activity flow terms (*p*_FWE_ < .05, 718 comparisons).

**SFigure 4.**
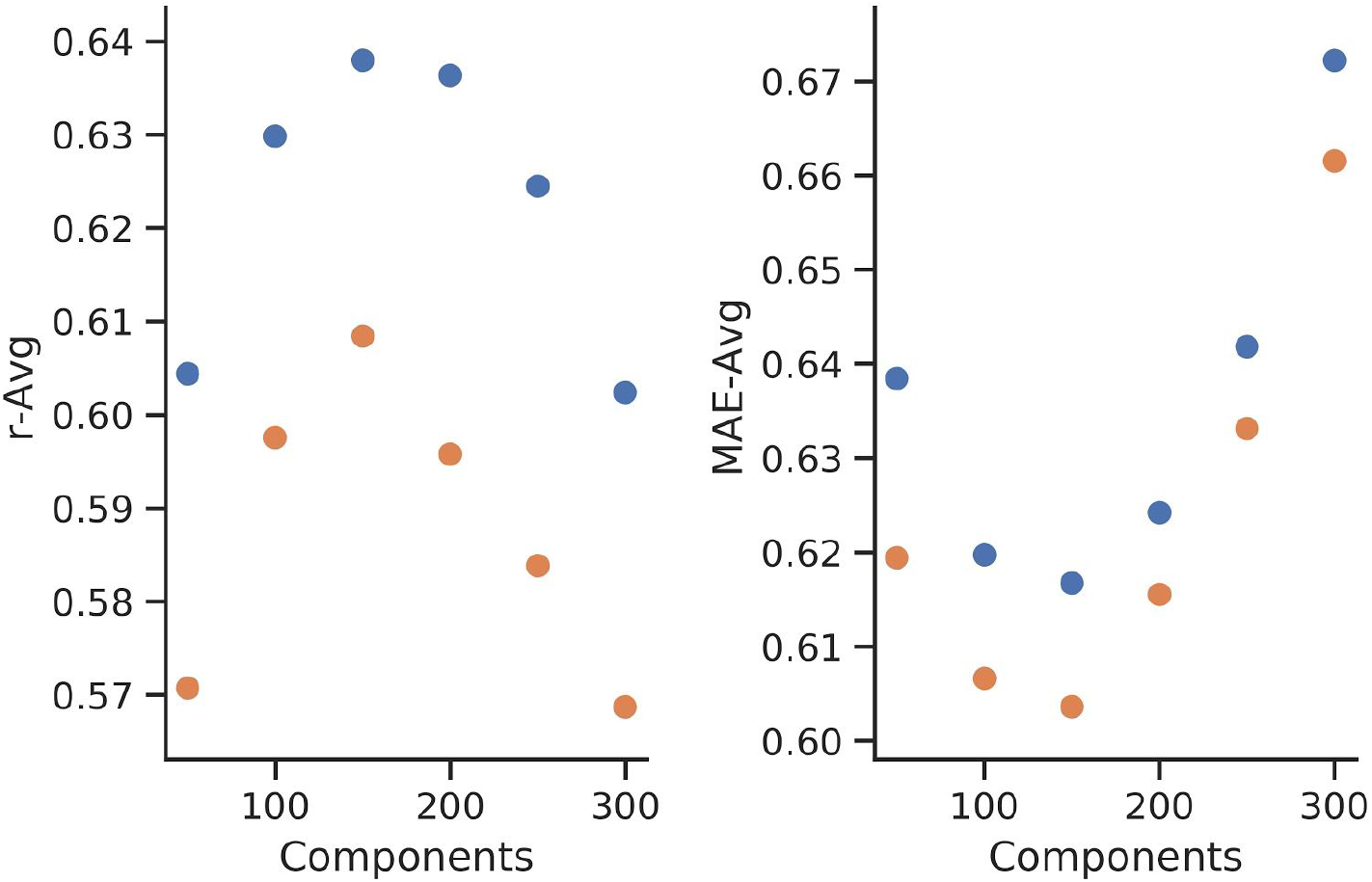
Effect of PCA regression component numbers in FC estimation on activity flow performance. Average activity flow performance when comparing the correlation with real data (*y*-axis, r-Avg, left) and the mean absolute error (*y*-axis, MAE-Avg, right) for healthy control (blue data) and schizophrenia cohorts (orange data). Multiple components were contrasted (*x*-axis) with peak performance at 150 components. Larger component sizes (e.g., 300) showed diminished performance, potentially due to overfitting. 100 components were used in the main analyses, which was selected apriori.

**SFigure 5.**
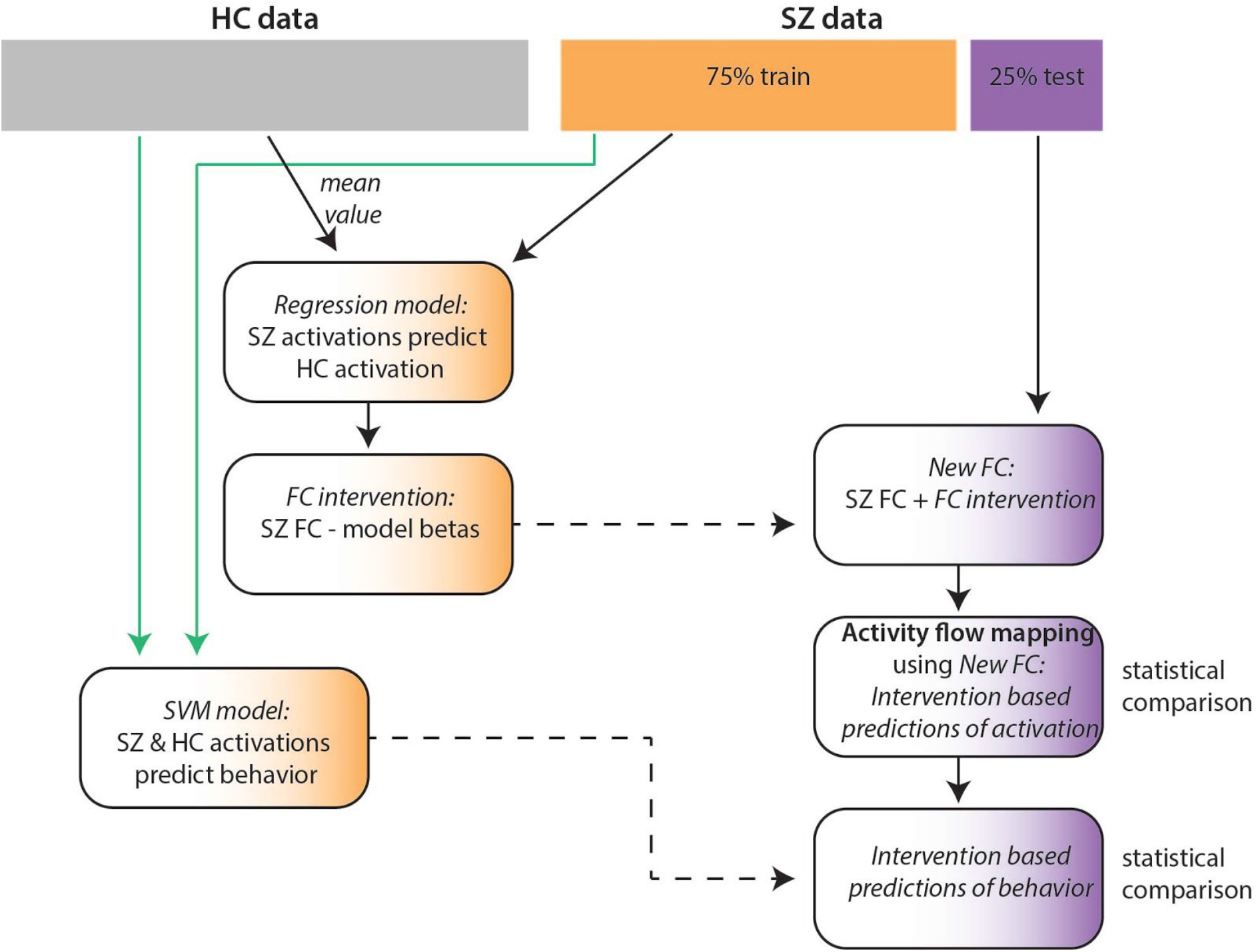
Schematic of hypothetical FC intervention fitting and testing procedure. To estimate the connectivity intervention a four fold cross validation scheme was used whereby models were fitted using the HC data and a subset of the SZ data (orange) and then tested on the held out SZ test cohort (purple).

**STable 1.** Correlation between region of interest averaged activity and behavioral measures

| Measure | Correlation (r) | $p_{FDR}$ | Significant |
| --- | --- | --- | --- |
| Spatial capacity working memory task | -0.37 | < 0.005 | TRUE |
| CA Verbal Learning: Short delay free recall | -0.27 | 0.03 | TRUE |
| CA Verbal Learning: Short delay cued recall | 0.22 | 0.15 | FALSE |
| CA Verbal Learning: Long delay free recall | 0.1 | 0.84 | FALSE |
| CA Verbal Learning: Long delay cued recall | -0.15 | 0.5 | FALSE |
| CA Verbal Learning: Long delay recognition | 0.02 | 0.99 | FALSE |
| Wechsler Memory: Symbol span | 0.04 | 0.99 | FALSE |
| Wechsler Memory: Visual reproduction immediate recall | 0 | 0.99 | FALSE |
| Wechsler Memory: Visual reproduction delayed recall | -0.04 | 0.99 | FALSE |
| Wechsler Memory: Visual reproduction recognition | -0.46 | < 0.005 | TRUE |
| Wechsler Memory: Digit span forward | 0.15 | 0.5 | FALSE |
| Wechsler Memory: Digit span backward | -0.37 | < 0.005 | TRUE |
| Wechsler Memory: Digit span sequencing | -0.32 | < 0.005 | TRUE |
| Wechsler Adult Intelligence: Matrix reasoning | -0.36 | < 0.005 | TRUE |
| Wechsler Adult Intelligence: Letter/number sequencing | -0.33 | < 0.005 | TRUE |
| Wechsler Adult Intelligence: Vocabulary | -0.32 | < 0.005 | TRUE |
| Color Trail: Interference index | -0.27 | 0.03 | TRUE |
| Task Switching: Accuracy | -0.19 | 0.28 | FALSE |
| Task Switching: Interference | -0.17 | 0.42 | FALSE |
| Task Switching: Switching cost | -0.23 | 0.11 | FALSE |
| Task Switching: Residual switching cost | -0.34 | < 0.005 | TRUE |
| Attention Network Task: Interference RT | 0.04 | 0.99 | FALSE |
| Go/No Go: Hit rate | -0.2 | 0.27 | FALSE |
| Go/No Go: Hits median RT | -0.24 | 0.11 | FALSE |
| Go/No Go: False alarm rate | -0.2 | 0.27 | FALSE |
| Delis-Kaplan Executive Function: English verbal fluency | -0.23 | 0.11 | FALSE |

Note: full details regarding the out of scanner measures in the CNP dataset can be found in Kebets et al., (2019).

**STable 2.** Medication use in the SZ cohort

|  |  |
| --- | --- |
| Current Medication | 31 Antipsychotic |
|  | 14 Antidepressant |
|  | 11 SHA |
|  | 7 Mood stabilizer |
|  | 5 Other |
|  | 1 Stimulant |

Note: Medication classes comprised antipsychotics, mood stabilizers, antidepressants, sedatives/hypnotics/anxiolytics (SHA), stimulants, and 'other'. A subset of participants were medicated with multiple drug classes simultaneously. Full details can be found in the original CNP data paper (Kebets et al., 2019; Poldrack et al., 2016).

